# Discovery of optimal cell type classification marker genes from single cell RNA sequencing data

**DOI:** 10.1101/2024.04.22.590194

**Authors:** Angela Liu, Beverly Peng, Ajith V. Pankajam, Thu Elizabeth Duong, Gloria Pryhuber, Richard H. Scheuermann, Yun Zhang

## Abstract

The use of single cell/nucleus RNA sequencing (scRNA-seq) technologies that quantitively describe cell transcriptional phenotypes is revolutionizing our understanding of cell biology, leading to new insights in cell type identification, disease mechanisms, and drug development. The tremendous growth in scRNA-seq data has posed new challenges in efficiently characterizing data-driven cell types and identifying quantifiable marker genes for cell type classification. The use of machine learning and explainable artificial intelligence has emerged as an effective approach to study large-scale scRNA-seq data. NS-Forest is a random forest machine learning-based algorithm that aims to provide a scalable data-driven solution to identify minimum combinations of necessary and sufficient marker genes that capture cell type identity with maximum classification accuracy. Here, we describe the latest version, NS-Forest version 4.0 and its companion Python package (https://github.com/JCVenterInstitute/NSForest), with several enhancements to select marker gene combinations that exhibit highly selective expression patterns among closely related cell types and more efficiently perform marker gene selection for large-scale scRNA-seq data atlases with millions of cells. By modularizing the final decision tree step, NS-Forest v4.0 can be used to compare the performance of user-defined marker genes with the NS-Forest computationally-derived marker genes based on the decision tree classifiers. To quantify how well the identified markers exhibit the desired pattern of being exclusively expressed at high levels within their target cell types, we introduce the On-Target Fraction metric that ranges from 0 to 1, with a metric of 1 assigned to markers that are only expressed within their target cell types and not in cells of any other cell types. NS-Forest v4.0 outperforms previous versions on its ability to identify markers with higher On-Target Fraction values for closely related cell types and outperforms other marker gene selection approaches at classification with significantly higher F-beta scores when applied to datasets from three human organs - brain, kidney, and lung.

## Introduction

Single-cell/single-nucleus RNA sequencing (scRNA-seq) methods have become an established approach for measuring cell transcriptional phenotypes and better understanding distinct cell types and their states based on gene expression patterns. Cell types can be defined as distinct cell phenotypes that include both canonical cell types and discrete cell states^1^. Efforts to define and categorize these cell types using advanced single cell technologies have been ongoing over the past decade, including the Human Cell Atlas (HCA)^2^, the NIH Human BioMolecular Atlas Program (HuBMAP)^3^, and the NIH BRAIN Initiative^4^. These efforts have led to consortium-scale datasets from multiple tissues/organs across the human body. For example, an early BRAIN Initiative study of the middle temporal gyrus (MTG) region in the human brain identified 75 distinct brain cell types with a dataset of approximately 16,000 nuclei^5^. A recent study on the transcriptomic diversity across the whole human brain revealed 461 cell type clusters and 3313 subclusters, with a final dataset comprised of more than three million cells^6^. The HuBMAP consortium covers other major human organs, including the kidney^7^ and lung^8^, resulting in a collection of 898 cell types with approximately 280 million cells across multi-omics assays^9^.

While the number of cell types being identified in these scRNA-seq data atlases is increasing rapidly, there is a lack of a scalable and generalizable marker gene selection method that can systematically characterize these newly identified cell types for downstream use cases, such as for designing spatial transcriptomics gene panels and semantic representation of cell types in biomedical ontologies. Historically, two main approaches have been used to identify cell type-specific marker genes from scRNA-seq data: differential expression (DE) analysis, and manual curation of gene lists using prior domain knowledge. For example, the anatomical structures (AS), cell types (CT), and biomarkers (B) ASCT+B tables^10^ provided by the HuBMAP consortium use markers found in the scientific literature curated by domain experts for most of the organs in HuBMAP^11^. This approach is not only infeasible for large-scale datasets, but also leads to potentially incomplete (missing markers) or redundant (markers of parent cell type being used for child cell type) information for the most granular cell types. Alternatively, DE genes selected by modified Wilcoxon rank sum (performed using the “presto” R package) or related statistical tests are used in the popular Azimuth^12^ web application for those cell types represented in the references. The DE approach selects genes based on the gene expression distributions and the adjusted p-values produced by a chosen testing method, which does not directly test the ability to classify cell types. Therefore, we formally introduce the notion of “cell type classification marker gene combinations” for scRNA-seq data, which must meet the following criteria: 1) each gene is expressed in the majority of cells of a given type, 2) each gene displays a “binary expression pattern” (i.e., highly expressed in the target cell type and little to no expression in other cell types), 3) gene combinations are optimized for cell type classification using metrics that quantify classification confidence, and 4) combinations can be reproduced by a generalizable method.

For the above-described challenge, we have proposed to use a machine learning approach to identify marker genes for cell type classification from scRNA-seq data and developed the NS-Forest method^13,14^. NS-Forest uses the random forest machine learning algorithm to select informative gene features (or markers) that are optimized for cell type classification. Random forest is a machine learning classification model that is well-known for retaining high explainability, which is preferable for biomedical use cases.

NS-Forest was first introduced in 2018 as an algorithm that takes in scRNA-seq data and outputs the minimum combination of necessary and sufficient features that capture cell type identity and uniquely characterize a discrete cell phenotype^1^. In NS-Forest v1.3^14^ (the first publicly released version), the method first produces a list of top gene features (marker candidates) for each cell type ranked by Gini index calculated in the random forest model. (Figure 1 summarizes major steps of the NS-Forest workflow compared across all versions.) The median gene expression value of each potential marker within the target cell type is calculated as the expression threshold to determine the number of true/false positives/negatives for each marker candidate in each cell type. Finally, the minimum set of markers for each cell type is determined by evaluating the unweighted F1-score following the stepwise addition of each of the ranked genes for each cell type.

**Figure 1.**
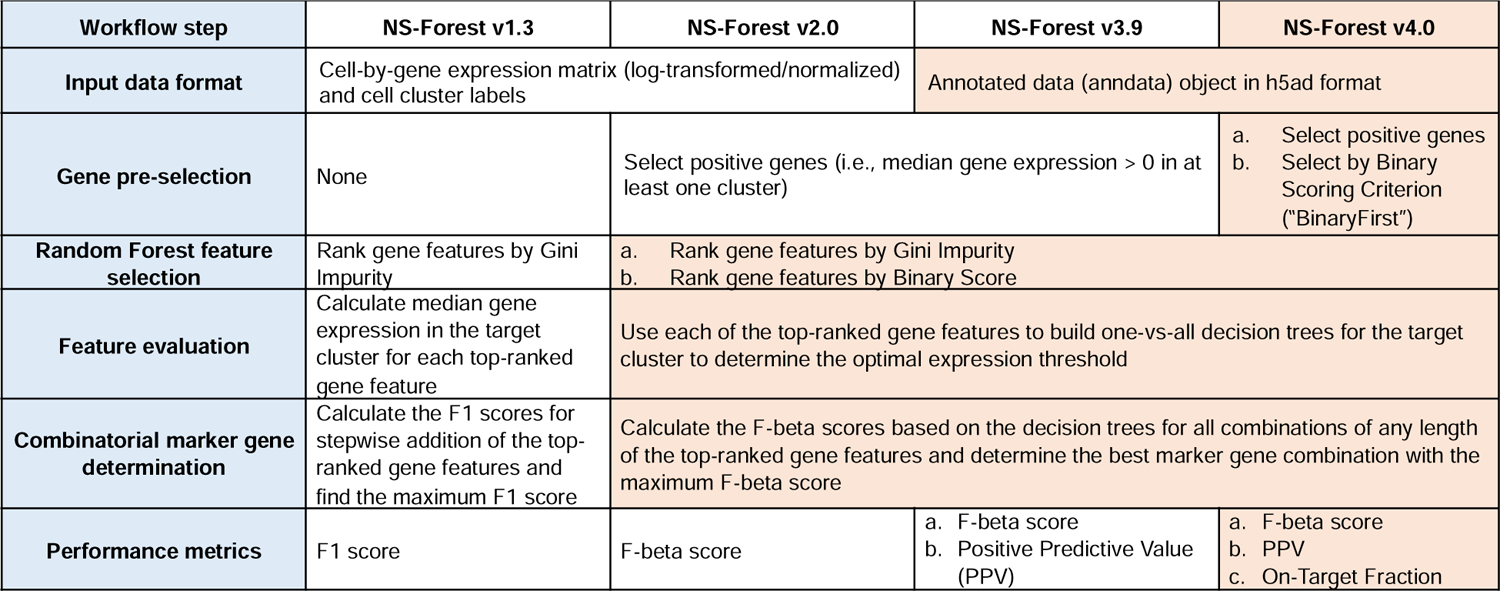
Major steps of the NS-Forest workflow compared across all versions.

NS-Forest v2.0^13^ was developed in 2021, and introduced the concept of the Binary Expression Score, a metric used to quantify how well a marker gene exhibits a “binary expression pattern” in which the marker gene is expressed at high levels in the majority of cells of the target cell type and not in cells of other cell types. Version 2.0 uses Binary Expression Score as a post random forest ranking step to preferentially select genes with the desired binary expression pattern, in addition to filtering out genes with negative expression levels, after the initial feature selection process from the random forest classifier. Instead of simply using each gene’s median expression value within the target cluster to determine its expression threshold, version 2.0 builds a one-versus-all decision tree for each marker candidate to derive the optimal expression level for classification. Finally, the F-beta score is calculated for all possible combinations of the top-ranked, most binary-expressed marker candidates in order to identify the best combination of markers with the maximum F-beta score. The F-beta score is different from the F1 score in that it is the weighted harmonic mean of the precision and recall (instead of just the harmonic mean), with the beta parameter weight adjustment allowing for emphasis of either precision or recall. In version 2.0 and all following versions, beta is set to 0.5 by default to weight precision higher than recall, to control for excess false negative values introduced by the dropout technical artifact in scRNA-seq experiments.

NS-Forest v3.9 is algorithmically very similar to version 2.0, and mainly differs by the format of the data used as input to the algorithm. Instead of the simple cell-by-gene expression matrix, where each entry contains the log-transformed or normalized expression level of each gene in each cell along with the cluster labels, version 3.9 takes the annotated data (anndata)^15^ object in the .h5ad file format as input. Version 3.9 also provides calculation of the Positive Predictive Value (PPV) metric (precision) for quantifying the classification performance of the algorithm in addition to the F-beta score, emphasizing the pragmatic importance of the predicted positives in many of the applications.

One of the observed lingering weaknesses in versions 2.0 and 3.9 was the lower performance of NS-Forest marker genes in distinguishing between closely-related cell types with similar transcriptional profiles. In the human brain middle temporal gyrus (MTG) dataset^5^ that was used to develop previous versions of NS-Forest, there exist several of these closely-related cell type groups, especially within the VIP, PVALB, and L4 neuronal cell subclasses (see Results section). Here, we describe NS-Forest v4.0, which adds an enhanced feature selection step to improve discrimination between similar cell types without sacrificing the overall classification performance. This new “BinaryFirst” step enriches for candidate genes that exhibit the binary expression pattern as a feature selection approach prior to the random forest classification step. The BinaryFirst strategy effectively reduces the complexity of the input feature set for the random forest classifier, allowing for the preferential selection of informative binary markers during the iterative random forest process and thus resulting in a more distinct set of marker genes.

## Results

### Informative gene selection prior to random forest in NS-Forest version 4.0

The most significant change to the workflow of NS-Forest in version 4.0 is the introduction of the BinaryFirst module that is implemented in the gene pre-selection step of the workflow (Figure 1). The BinaryFirst strategy is designed to enrich for candidate genes that exhibit the desired gene expression pattern prior to the random forest feature ranking. This step pre-selects gene candidates that have a Binary Expression Score value that is greater than or equal to a dataset-specific threshold based on the distribution of the Binary Expression Scores of all genes in the dataset (Figure 2). In version 4.0, four threshold options used in the BinaryFirst step were implemented: ‘none’, ‘BinaryFirst_mild’, ‘BinaryFirst_moderate’, or ‘BinaryFirst_high’ (see Methods). If the threshold is ‘none’, the algorithm is the same as version 3.9. The other thresholds are calculated based on the distribution of Binary Expression Scores from all genes to account for the dataset-specific gene expression variabilities arising from many factors, including the organ/tissue type, sample pre-processing, sequencing platform, etc. As the thresholding values increase, the number of selected genes decrease. Thus, the BinaryFirst strategy effectively reduces the feature space that the random forest classifier must search over in the subsequent workflow step of NS-Forest and serves as informative dimensionality reduction. An scRNA-seq dataset typically contains tens of thousands of genes, but the majority will not be useful as marker genes for any given cell type clusters. Random forest is an ensemble machine learning method that uses the bagging technique to train a large number of decision trees. The bagging technique is an iterative procedure of random selection of features and, therefore, is time-consuming for classifying cell types from scRNA-seq data^16^. Each of the decision trees in the forest is constructed of nodes, at which the data is split into groups that optimize class purity using randomly selected features. Because performing an exhaustive search of all possible combinations of features at each split in the decision trees is computationally intractable, random forest classifiers can produce sub-optimal collections of decisions trees due to the random selection of available features^17^. In previous versions of NS-Forest, the large number of genes in the input datasets reduced the likelihood that the optimal genes for classification would be adequately sampled. Our hypothesis was that by reducing the size of the set of gene candidates, BinaryFirst would be able to adequately sample all of the candidate input genes with a reasonable number of decision trees, thereby simplifying the task of the random forest classifier, while simultaneously reducing the overall runtime of NS-Forest. In summary, NS-Forest v4.0 utilizes the BinaryFirst strategy to enhance the stability and classification performance of the random forest classifier by pre-selecting informative features from scRNA-seq data.

**Figure 2.**
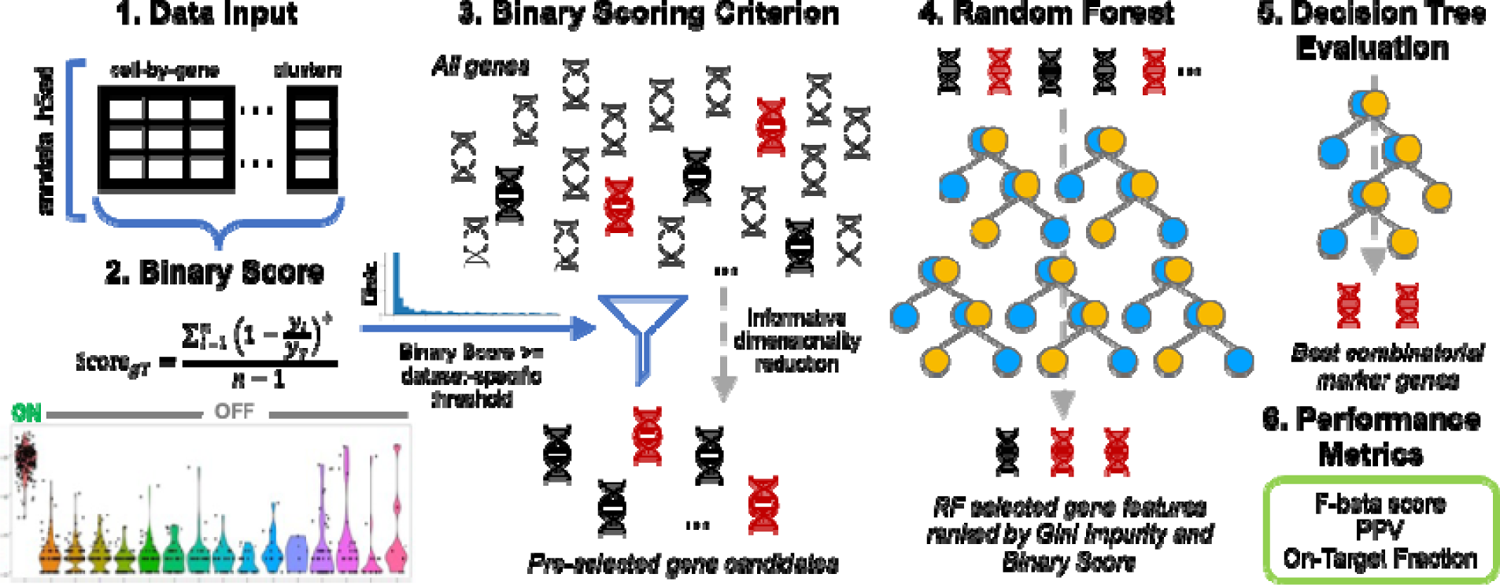
NS-Forest version 4.0 workflow. The algorithm uses an anndata object in .h5ad format, containing the cell-by-gene expression matrix and cluster labels for each cell, as data input (step 1). The median gene expression for each gene in each cluster (i.e., a cluster-by-gene median matrix) is calculated and genes that have positive median expression in at least one cluster are pre-selected (not shown). The Binary Expression Score is then calculated for each cluster-gene pair (step 2) producing a cluster-by-gene Binary Score matrix (note that a gene may have different Binary Score values in different clusters), and a dataset-specific threshold is calculated based on the Binary Score distribution and user-selected mild, moderate, or high criterion. This threshold value is used to select candidate genes for each cluster with a Binary Expression Score greater than or equal to the threshold (step 3). These candidate genes are passed to build binary classification models for each cluster using the random forest (RF) machine learning method. Features (genes) are extracted from the RF model and ranked by the Gini Impurity index, and the top RF features are then reranked by their pre-calculated Binary Scores (step 4). A short list of the top-ranked candidate genes that are not only ranked high in the RF classification models but also have high Binary Scores are passed for decision tree feature evaluation and determining the best marker gene combination. A single-split decision tree is built for each evaluated gene for determining the optimal expression threshold for classification. All combinations of any length of these genes are considered using ‘AND’ logic to combine the decision trees, and the best combination is determined by the highest F-beta score as an objective function for optimizing the overall classification performance (step 5). The F-beta score, Positive Predictive Value (PPV), On-Target Fraction, as well as true/false positive/negative classification values are reported for each cluster, serving as metrics for evaluating the performance of the final maker gene combinations (step 6).

### Improved marker gene selection on the human brain dataset

The performance of NS-Forest v4.0 was assessed on the same human middle temporal gyrus (MTG) brain dataset that was used to evaluate previous versions of NS-Forest^5^. To determine the best thresholding criterion, the mild, moderate, and high BinaryFirst configurations were compared for the version 4.0 algorithm. The evaluation is based on the On-Target Fraction metric (see Methods), which is specifically designed to quantify how much of the marker gene expression is restricted to the target cluster. Comparing these three configurations, the On-Target Fractions significantly increase as the stringency of BinaryFirst thresholding increases (Figure 3A), which results in fewer candidate genes for random forest construction (Figure 3B). This suggests that the gene pre-selection step helps the NS-Forest algorithm better select on-target marker genes while effectively reducing the dimensionality of the input gene space. Hereinafter, the ‘BinaryFirst_high’ configuration will be the default setting for NS-Forest v4.0, unless specified otherwise.

**Figure 3.**
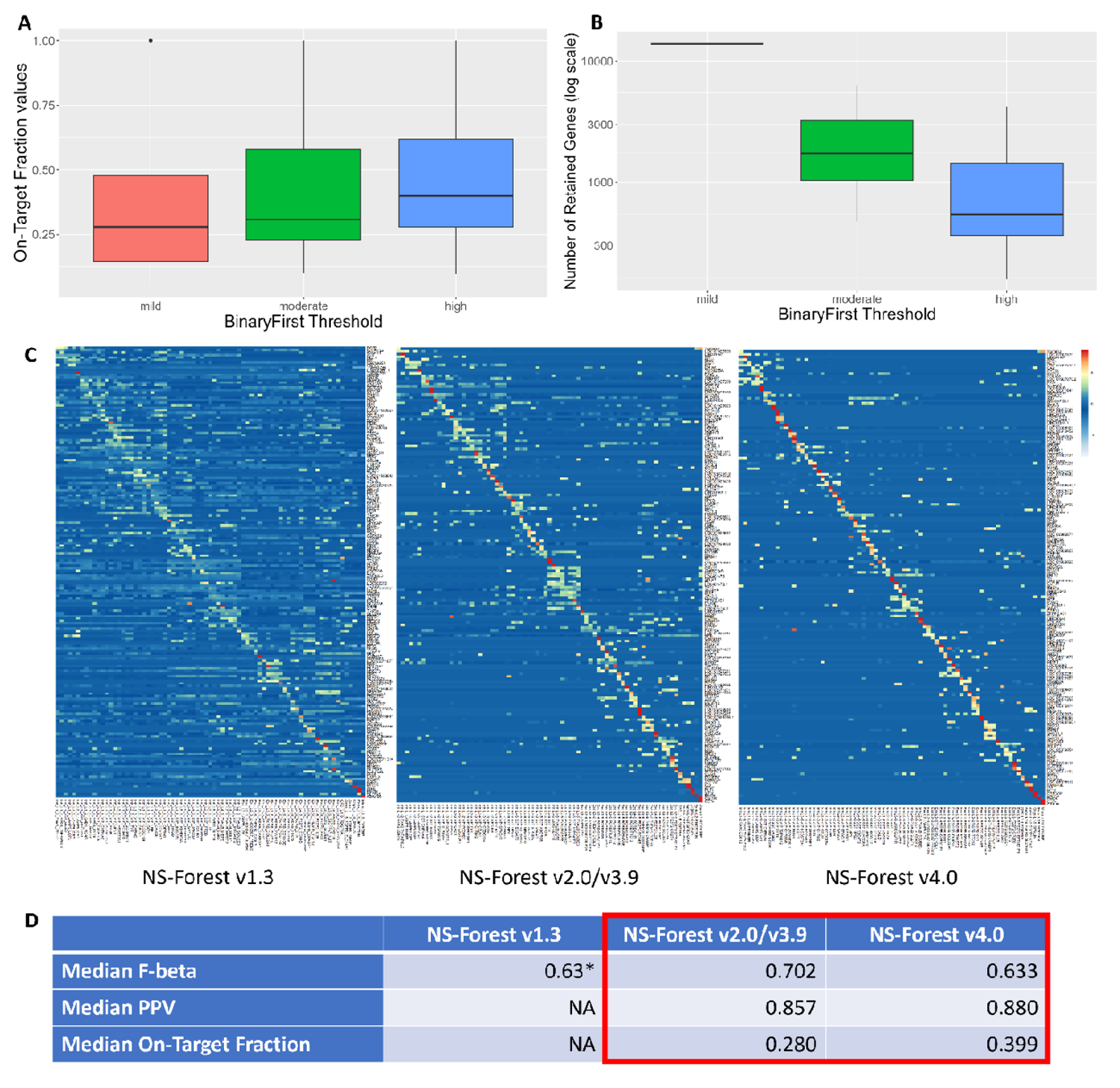
NS-Forest performance evaluated on the human MTG dataset. **(A)** Boxplots displaying the distribution of On-Target Fraction values across the 75 clusters in the human MTG dataset from running NS-Forest v4.0 with mild, moderate, and high BinaryFirst configurations. Paired t-test results: mild vs. moderate p-value = 0.01, moderate vs. high p-value = 1.79e-04, mild vs. high p-value = 1.68e-06. **(B)** Boxplots displaying the distribution of the number of genes retained after the BinaryFirst thresholding step across the 75 clusters from the human MTG dataset from running NS-Forest v4.0 with mild, moderate, and high BinaryFirst configurations. **(C)** Heatmaps of NS-Forest v1.3, v2.0/v3.9, and v4.0 marker genes for the 75 cell type clusters from the human MTG dataset. The colors correspond to the normalized median expression level (log2-transformed counts per million) for the marker gene (rows) in a given cell type cluster (columns), with high expression in red/yellow, and low expression in blue/white. The clusters are ordered according to the hierarchical dendrogram provided in the original study (Figure 1c in Hodge et al. 2019). **(D)** Comparison of the performance metrics of the corresponding versions of NS-Forest results shown in (C). *Note that the unweighted F1 score is used in v1.3. The On-Target Fraction difference between v2.0/3.9 and v4.0 corresponds to the mild and high BinaryFirst threshold comparison in (A).

The performance of all versions of NS-Forest were compared on the same brain dataset. The overall performances of different versions are shown in Figure 3C, where a noticeable decrease in off-target expression is observed across the three heatmaps from versions 1.3 to 2.0/3.9 to 4.0. In the most ideal scenario where there exist marker genes that are exclusively expressed for each cluster, marker gene expression would only be observed in a stair-step pattern along the diagonal axis in such expression heatmaps. By this standard, NS-Forest v4.0 shows the cleanest diagonal pattern in its heatmap. Improvement is also observed when comparing the performance metrics, as the median PPV and the median On-Target Fraction increased from v2.0/v3.9 to v4.0 across all 75 clusters (Figure 3D). The improvement in PPV (precision) means that version 4.0 does a better job at identifying marker genes that, when used as features by decision tree classifiers, lead to improved performance at identifying cells that belong to each unique cell type by reducing the number of false positives in this classification task. The increase in On-Target Fraction further supports this claim that version 4.0 is able to identify marker genes that are exclusively expressed at high levels in their target clusters. This improvement in the PPV and On-Target Fraction is slightly offset by a small decrease in the median F-beta score between the two versions. The tradeoff in the F-beta score was previously discussed in Aevermann et al.^13^ when comparing the off-the-shelf random forest marker candidates strictly ranked by feature importance (Gini Impurity) to the top candidates re-ranked by their Binary Expression Score values. Aevermann et al. demonstrated that the marker genes selected with significantly higher Binary Expression Scores are more useful for many downstream assays such as RT-PCR and spatial transcriptomics. A similar trend was observed in this current comparison of versions 2.0/3.9 and version 4.0: the Binary Expression Scores are higher for v4.0 than for v2.0/3.9, with an average of 0.971 compared to 0.936. This is consistent with the observations of a cleaner diagonal pattern in the heatmap and higher On-Target Fraction values for version 4.0.

In total, NS-Forest versions 2.0/3.9 and version 4.0 identified 168 and 167 total marker genes, and 154 and 147 unique markers, respectively, to optimally distinguish between the 75 cell type clusters in the human MTG dataset. 85 of the total markers (∼51% of the markers identified by v2.0/3.9) were identified for the same cluster for both sets. The full list of NS-Forest v4.0 marker genes on the human MTG dataset are available in Supplementary Table 1.

### Localized improvement in marker gene specificity for closely related cell types

One of the motivations for developing NS-Forest v4.0 was to improve the previously sub-optimal performance on specific subclades of closely related cell types in the MTG dataset that may be more difficult to distinguish compared to all other cell types. The MTG study found that the inhibitory neuron types are highly diverse but mostly sparse (45 types and 4,297 nuclei), and the excitatory neuron types span brain layers and are most similar to types in the same or adjacent layers (24 types and 10,708 nuclei)^5^. These highly similar yet distinct cell types are usually grouped as subclades in the hierarchical dendrogram (Supplementary Figure 1A). The performance of version 4.0 on the VIP (vasoactive intestinal polypeptide-expressing inhibitory neurons), PVALB (parvalbumin-expressing inhibitory neurons) and L4 (layer 4 excitatory neurons) subclades was examined using the three different BinaryFirst thresholds to determine if NS-Forest v4.0 would produce higher On-Target Fractions. Visually, there appears to be a clear improvement in the VIP subclade, as the amount of off-target expression (represented by the number of yellow squares not on the diagonal axis in the highlighted VIP box) looks to be substantially fewer in the heatmap for the ‘BinaryFirst_high’ configuration (Supplementary Figure 1B). The amount of on-target expression (represented by the amount of red and orange squares on the diagonal axis) appears to be greatest in this heatmap as well. While the pattern of the PVALB and L4 subclades is less obvious, the pattern of increased on-target expression and decreased off-target expression is still observable. These trends are also observed in the median On-Target Fraction values, as this value is the highest for all three subclades using the ‘BinaryFirst_high’ configuration (Supplementary Figure 1C).

The distribution of On-Target Fractions for each of the three subclades is consistent with these visual patterns. The boxplot showing the On-Target Fraction values for the ‘BinaryFirst_high’ threshold is clearly higher than that for the ‘BinaryFirst_mild’ and ‘BinaryFirst_moderate’ thresholds in all three subclades (Supplementary Figure 2). The p-value for comparing the mild and high BinaryFirst thresholds for the VIP subclade is statistically significant (p-value = 0.01), but the p-values for the L4 subclade and the PVALB subclade are not significant (p-value = 0.37 and 0.07, respectively); this is likely due to the small sample size for the paired t-test (the L4 subclade has 5 distinct cell types and the PVALB subclade has 6, whereas the VIP subclade has 21).

### Validation on additional datasets of human kidney and lung

Datasets from two other human organs - the human kidney dataset from the Kidney Precision Medicine Project (KPMP)^7^ and the human lung dataset from the Lung Airways and Parenchymal Map (LAPMAP)^8^, both contributing to the HuBMAP consortium – were used to validate the NS-Forest v4.0 performance. In the lung dataset, three annotation levels were evaluated: level 3 (L3), level 4 (L4), and level 5 (L5) subclasses. The kidney dataset has 75 distinct cell types, and the lung dataset has 61 cell types at the L5 subclass. For both the kidney and lung datasets, the heatmaps show more specific expression along the main diagonal with version 4.0 (Figure 4A), which complement the observed high On-Target Fraction values for these datasets (Figure 4B). These values are higher than for the brain dataset because the human brain is a more complex organ in terms of the diversity of related cell types. Version 2.0/3.9 already performs quite well at selecting marker genes for these two organs compared to the brain, and hence the improvement of version 4.0 is less obvious.

**Figure 4.**
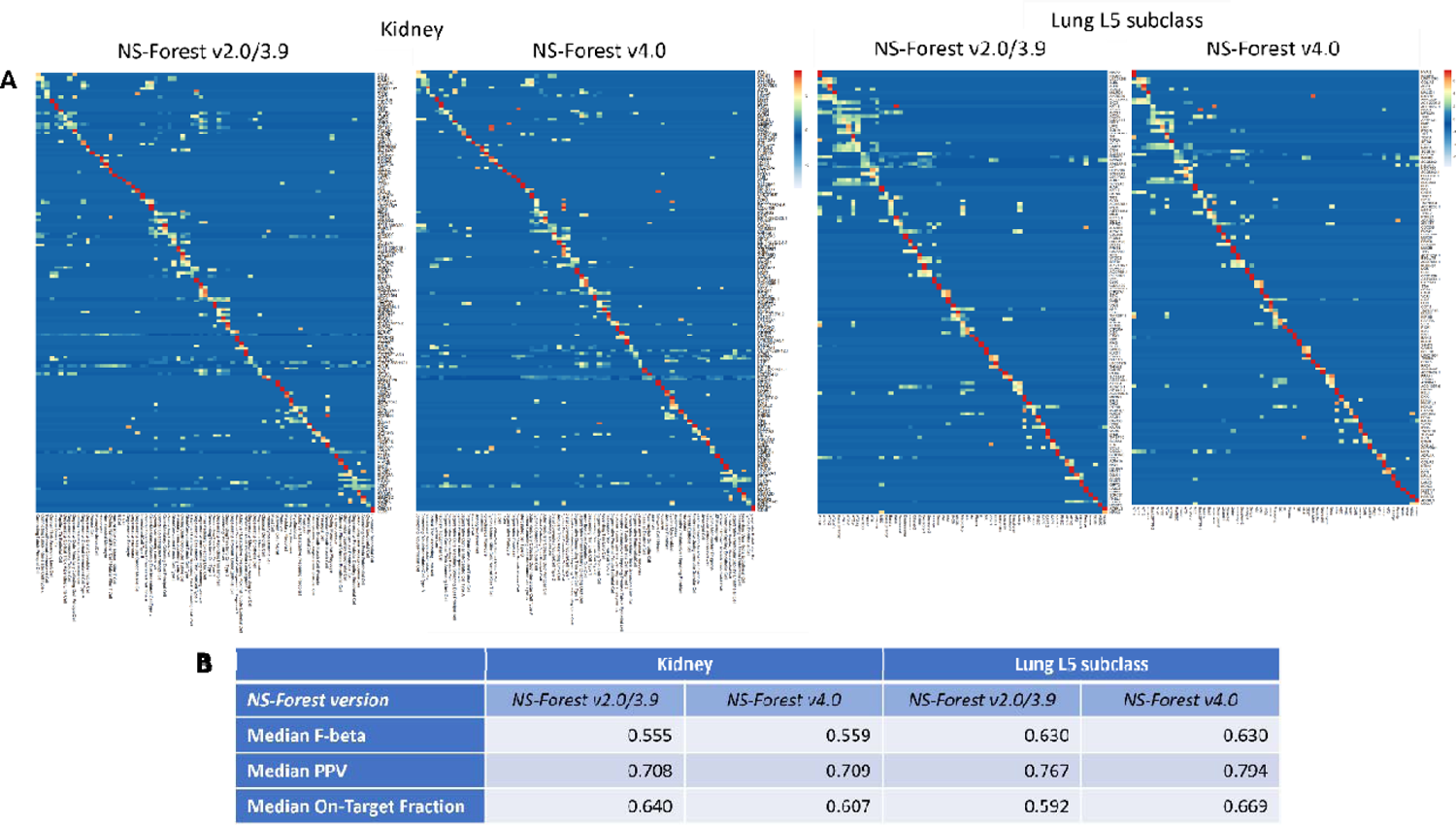
NS-Forest performance evaluated on other human organs (kidney and lung). **(A)** Heatmaps of NS-Forest v2.0/v3.9 and v4.0 marker genes for the 75 cell type clusters from human kidney dataset and 61 cell types from human lung L5 subclass dataset. The colors correspond to the normalized median expression level (log2-transformed counts per million) for the marker gene (rows) in a given cell type cluster (columns), with high expression in red/yellow, and low expression in blue/white. The clusters are ordered according to the hierarchical ordering in the dendrogram generated by the scanpy package (*scanpy.tl.dendrogram*) using default settings. **(B)** Comparison of the performance metrics corresponding to the NS-Forest results shown in (A).

Comparing versions 2.0/3.9 and 4.0 (Figure 4B), the median F-beta scores are very similar in both versions for each organ. In the kidney dataset, a slight improvement in PPV and a slight decrease in On-Target Fraction were observed going from version 2.0/3.9 to version 4.0. It is interesting to note that although the On-Target Fraction is slightly lower, the number of false positive classifications is much fewer in version 4.0 (dropped from an average of 312.5 cells in version 2.0/3.9 to 198 cells in version 4.0), confirming that the version 4.0 marker genes are selected for optimal classification. In the lung L5 subclass dataset, increases in both PPV and On-Target Fraction were observed with version 4.0.

NS-Forest v2.0/3.9 and v4.0 identified 157 and 169 total marker genes for the kidney dataset, and 144 and 151 unique markers, respectively, to optimally distinguish between the 75 cell types (Supplementary Tables 2-3). 117 of the total markers (75% of the markers identified by v2.0/3.9) were identified for the same cluster for both sets. NS-Forest v2.0/3.9 and v4.0 identified 131 and 126 total marker genes for the lung L5 dataset, and 125 and 121 unique markers, respectively, to optimally distinguish between the 61 cell types. 107 of the total markers (82% of the markers identified by v2.0/3.9) were identified for the same cluster for both sets. Similar results were obtained running NS-Forest on the L4 and L3 subclasses (49 and 44 types, respectively) of the same lung dataset (Supplementary Figure 3 and Supplementary Tables 4-5), suggesting that it is easier to select marker genes at less granular levels.

### Improvement in runtime in version 4.0

In addition to the improvements in the classification performance, the overall runtime of NS-Forest v4.0 (by default, ‘BinaryFirst_high’ is used) is much lower than that of v2.0/3.9 in all three human organ datasets (Table 1), with the ratio of runtime (v4.0 to v2.0/3.9) ranging from 0.10 to 0.26 across the datasets. In all three datasets, the ‘BinaryFirst_mild’ configuration did not filter out any genes, indicating that more than half of the genes have a value of 0 for their Binary Expression Score and thus, a value of 0 for their median expression per cluster. We note that for all three datasets, there is a large decrease in runtime going from v2.0/3.9 to v4.0 with the ‘BinaryFirst_moderate’ configuration, and a less substantial decrease going from the moderate to ‘BinaryFirst_high’ configuration. These differences in runtime correspond to the decreases in the average number of genes left per distinct cell type after the BinaryFirst step. When the high threshold was used, 1-7% of the total original genes passed the BinaryFirst threshold in the three datasets. This demonstrates that the majority of genes in these datasets have low Binary Expression Scores, and that the distribution of Binary Expression Scores in these datasets is heavily right-skewed (Supplementary Figure 4), which is generally true for all scRNA-seq data. Overall, the BinaryFirst step can dramatically reduce the number of candidate genes that are considered as potential markers as input to the random forest step and simultaneously provide improvement in important measures of classification performance.

**Table 1.**
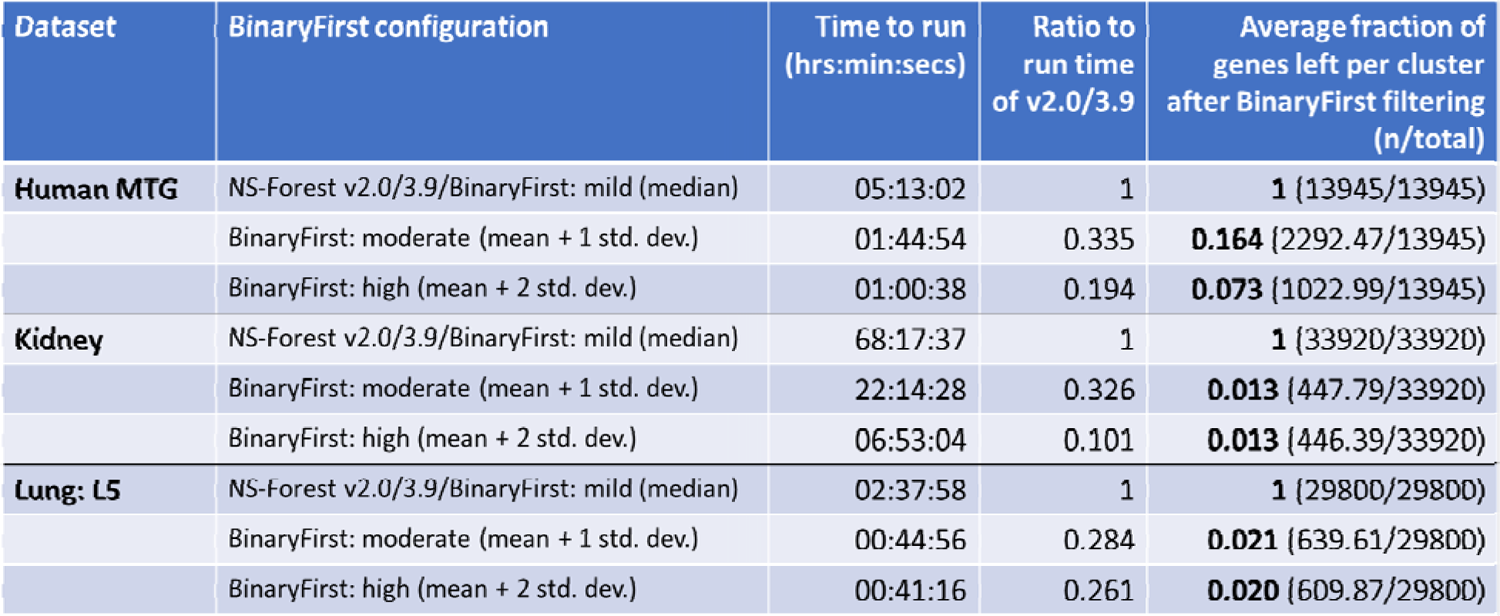
Comparison of runtime and number of genes that passed the BinaryFirst filtering criterion between different BinaryFirst configurations.

### Marker gene comparison for the Human Lung Cell Atlas

While the goal of NS-Forest is to define the minimum set of necessary and sufficient marker genes to classify cell types from scRNA-seq data, other popular marker gene approaches aim to define marker genes by identifying genes that are differentially expressed between cell types (e.g., Azimuth^12^), or by manually curating knowledge historically reported in the scientific literature (e.g., ASCT+B from the HuBMAP consortium^10^). Though Azimuth and ASCT+B provide pan-organ marker gene lists as centralized resources, it is more often that individual studies provide their own marker gene lists derived from each specific dataset. To compare the performance of different marker gene selection approaches, the Human Lung Cell Atlas (HLCA) core dataset was used as a common comprehensive data resource, consisting of ∼0.5 million cells clustered into 61 cell types from healthy lung tissues from 107 individuals^18^. The HLCA authors provided cell type-specific marker genes by iteratively subsetting the atlas into sequentially granular classifications and filtering for unique genes within the compartments (hereinafter, HLCA markers). Meanwhile, the lung single cell community also constructed the LungMAP single-cell reference (CellRef) to provide integrated information for both human and mouse lungs^19^. For this comparison, we consider five marker gene lists for healthy human lung cell types: NS-Forest markers, HLCA markers, CellRef markers, ASCT+B markers, and Azimuth markers. The HLCA markers are available in Supplementary Table 6 from the publication^18^. The CellRef markers were extracted from the human LungMAP CellCards, which are available from Supplementary Data 2 from the publication^19^. The ASCT+B markers are available from the “lung v1.4” table at the Human Reference Atlas portal (https://humanatlas.io/asctb-tables). The Azimuth markers derived from the HLCA core dataset are pre-calculated and available under “Human - Lung v2 (HLCA)” at the Azimuth portal (https://azimuth.hubmapconsortium.org/). Because CellRef and ASCT+B markers are not directly derived for the HLCA cell types, the Simple Standard for Sharing Ontological Mappings (SSSOM) guideline^20^ and Cell Ontology^21^ IDs were used to map the CellRef and ASCT+B cell types to the HLCA cell types, resulting in 33 and 18 exact matches, respectively.

To derive NS-Forest markers, NS-Forest v4.0 was applied to the HLCA core dataset and 122 marker genes (1-4 marker genes per type) for the 61 finest level cell types were identified (Supplementary Table 6). Figure 5A shows the NS-Forest marker gene expression in the 61 HLCA cell types ordered according to the dendrogram in Figure 5B. In this dendrogram, similar cell types are grouped according to the hierarchical clustering of the transcriptome profiles of these cell types; three major branches of immune cells, endothelial and stromal cells, and epithelial cells are observed. With the dendrogram ordering, the NS-Forest marker genes show a strong and clean expression pattern along the main diagonal in the expression dotplot (Figure 5A). Similar dotplots were produced for the other four marker gene lists (Supplementary Figure 5). The HLCA marker list contains 162 marker genes (1-5 markers per type) for the 61 cell types, which shows an expected diagonal pattern in the expression dotplot but with more off-diagonal expressions (Supplementary Figure 5A). The CellRef marker list contains 115 marker genes (2-7 markers per type) for the 33 exact matched cell types. The CellRef dotplot shows a relatively clean diagonal expresion pattern, although some genes show high levels of expression for multiple cell types (Supplementary Figure 5B). The ASCT+B marker list contains 80 marker genes (3-5 markers per type) for the 18 exact matched cell types. The ASCT+B dotplot is sparse (Supplementary Figure 5C) because the manual approach based on existing knowledge from the scientific literature does not capture the granularity obtained in single cell-resolution data. The Azimuth marker list contains 535 marker genes (8-10 markers per type) for 56 of these cell types (AT0, Hematopoietic stem cells, Hillock-like, Smooth muscle FAM83D+, and pre-TB secretory markers are not available). The Azimuth dotplot lacks a clean diagonal pattern with many of the genes being expressed at high levels across many similar cell types (Supplementary Figure 5D).

**Figure 5.**
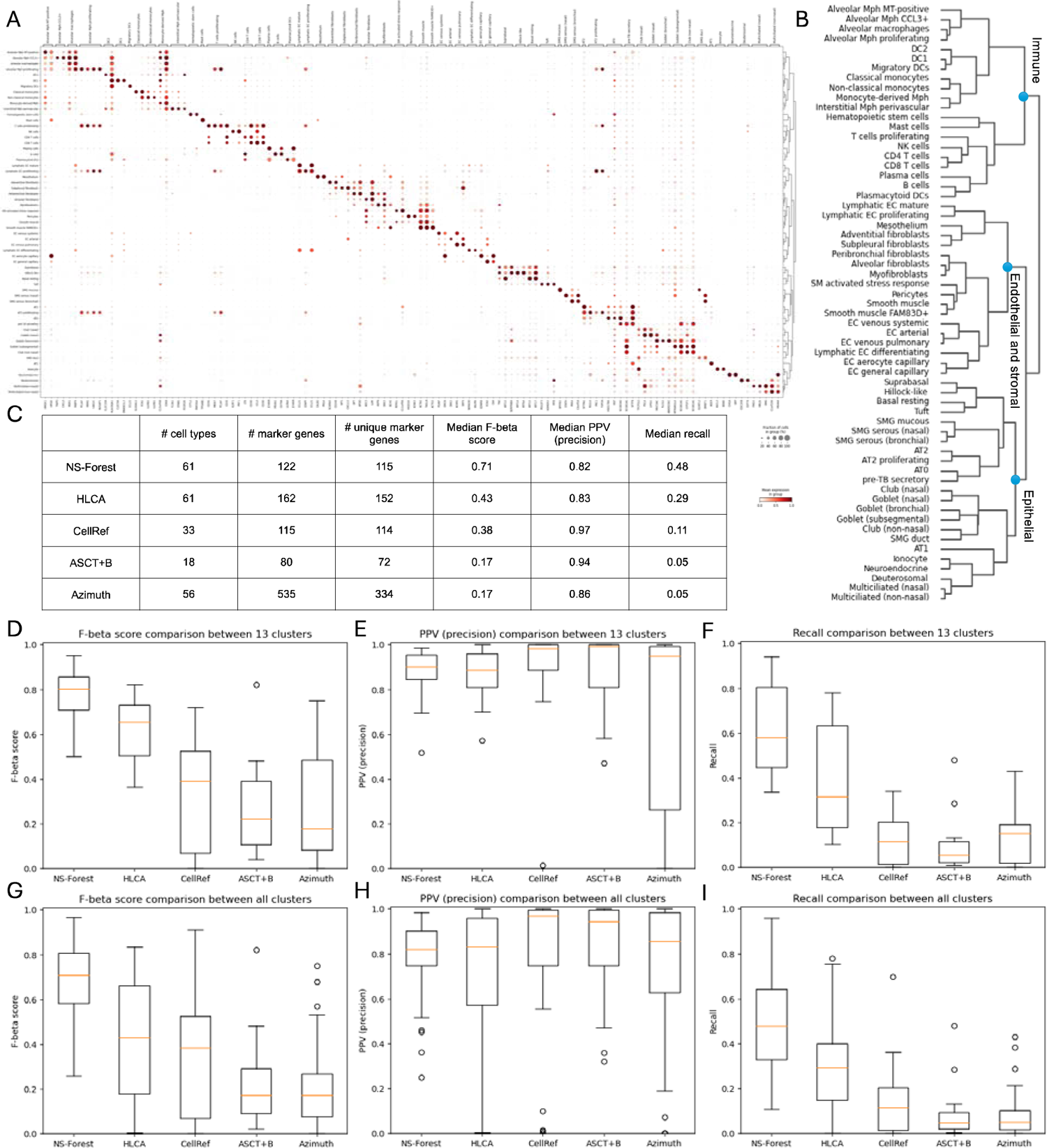
NS-Forest marker genes compared to other published HLCA marker gene lists. **(A)** Dotplot of the 122 NS-Forest marker genes on HLCA core cell types. **(B)** Dendrogram of 61 ann_finest_level cell types from HLCA core dataset corresponding to the rows in (A), generated before preprocessing. **(C)** Comparing number of cell types, number of markers, number of unique markers, median f-beta score, median PPV (precision), and median recall for the NS-Forest, HLCA, CellRef, ASCT+B, and Azimuth marker lists. **(D-F)** Boxplots of F-beta score, PPV (precision), and recall for the 13 cell types commonly characterized by all methods. **(G-I)** Boxplots of F-beta score, PPV (precision), and recall for all available cell types across methods.

The NS-Forest v4.0 Python package was also modularized to enable user-defined marker gene evaluation, allowing for direct comparison of the cell type classification metrics between different input marker lists. Using this approach, the performance of the five marker gene sets was compared using the HLCA data by calculating F-beta, PPV (precision), and recall for all cell types and directly comparing the medians (Figure 5C), along with the distributions of these performance metrics for the 13 cell types matched across all five reference datasets (Figure 5D-F) and for all matched cell types (Figure 5G-I). For the fair comparison of the 13 common cell types, the NS-Forest marker genes have the highest median F-beta score (0.80), followed by the HLCA markers (0.65), CellRef markers (0.39), ASCT+B markers (0.22), and Azimuth markers (0.18). Similar results were found using all cell types (NS-Forest: 0.71, HLCA: 0.43, CellRef: 0.38, ASCT+B: 0.17, and Azimuth: 0.17). The F-beta scores reflect the global patterns observed in the dotplots of these marker gene lists (Figure 5A and Supplementary Figure 5). The distributions of F-beta scores are significantly different between NS-Forest and all other methods (NS-Forest vs. HLCA: p = 0.0039, NS-Forest vs. CellRef: p = 2.0e-5, NS-Forest vs. ASCT+B: p = 2.1e-6, NS-Forest vs. Azimuth: p = 1.6e-5). Surprisingly, the median PPV (precision) scores are high for all five methods (0.89-0.99), with the ASCT+B being the highest, showing no significant difference between NS-Forest and the other methods (NS-Forest vs. HLCA: p = 0.70, NS-Forest vs. CellRef: p = 0.93, NS-Forest vs. ASCT+B: p = 0.81, and NS-Forest vs. Azimuth: p = 0.098). In contrast, the median recall values show a similar trend to the F-beta scores, with NS-Forest being the highest (0.58) and ASCT+B being the lowest (0.054) in the 13 common cell types. The distributions of recall values are significantly different between NS-Forest and all other methods (NS-Forest vs. HLCA: p = 0.036, NS-Forest vs. CellRef: p = 3.5e-7, NS-Forest vs. ASCT+B: p = 2.0e-6, and NS-Forest vs. Azimuth: p = 3.9e-6). Scatter plots for pairwise comparison between the NS-Forest markers and other markers for each cell type also show superior F-beta and recall results using NS-Forest marker combinations (Supplementary Figure 6), while having comparable performance in PPV (precision). Low F-beta and recall values for the other four methods are driven by excessive false negative predictions. Comparing all methods, NS-Forest produces the most comprehensive and concise list of marker genes with consistently higher F-beta scores for cell type classification.

## Discussion

This paper describes major algorithmic refinements made in NS-Forest v4.0 and its improved performance on the human brain MTG dataset used to develop the previous versions, as well as its performance on datasets from the human kidney and lung. The main motivation for developing version 4.0 was to improve the marker gene selection performance on closely related cell types by enriching for markers that exhibit the pattern of being highly and uniquely expressed in their target cell types without losing a significant amount of classification power. The BinaryFirst step was introduced in the NS-Forest workflow to enrich for candidate genes that exhibit the desired gene expression pattern. It is essentially an informative dimensionality reduction approach that effectively reduces the size of the set of candidate genes prior to the random forest classification step, which is usually the most time-consuming part of the algorithm. As a result, the overall runtime of the algorithm is substantially reduced. To explicitly demonstrate the improvements made by this algorithmic refinement, we introduced the On-Target Fraction metric that quantifies how well the NS-Forest marker genes are exclusively expressed in each distinct cell type.

Overall, this new version of NS-Forest demonstrated clear improvement in the human MTG brain dataset, which is the most complex organ evaluated in this study. Additional datasets representing the human kidney and lung were used to validate NS-Forest’s performance on data from other organs. NS-Forest v4.0 is now a comprehensive Python package that not only implements the algorithm for obtaining the marker genes, but also supports the marker gene evaluation functions in a machine learning framework for cell type classification. In this paper, we also presented a comparative analysis of the NS-Forest marker genes and other poplular marker gene lists. We formally established the notion of cell type classification marker genes, which are different from the notion of differentially expression genes. In the head-to-head comparison using the HLCA dataset with half a million cells, the NS-Forest marker genes showed superior performance than the HLCA, CellRef, ASCT+B and Azimuth marker sets.

NS-Forest marker genes can be used for multiple downstream experimental investigations, such as spatial transcriptomics gene panel design in the SpaceTx consortium^22^, and to produce marker gene sets designed to capture specific cell type properties. One of the main applications of NS-Forest identified marker genes is contributing to the definition of ontological classes of scRNA-seq data-driven cell types for incorporation into the official Cell Ontology^21^, as NS-Forest provides the minimum combinations of marker genes that can serve as a set of definitional characteristics of the cell types^21^. Such efforts have already begun as NS-Forest has contributed to the BRAIN Initiative Cell Census Network (BICCN) data ecosystem to derive the necessary and sufficient marker gene knowledge^23^. As such, the Provisional Cell Ontology (PCL) is generated in this manner for human, mouse, and marmoset primary motor cortex^24^. Among the general single cell community, there is a current lack of a formal, standardized representation of cell type clusters derived from the tremendous amount of scRNA-seq data and their transcriptional characterization that is widely accepted by the scientific community. One of the challenges associated with formalizing such a representation is the aspect of scaling up the semantic knowledge representations to keep up with the rate at which single cell transcriptomic data and analyses are being produced today. To this end, NS-Forest appears well-suited to help alleviate some of the challenges associated with such a task, especially with the enhancements introduced in version 4.0.

## Methods

### BinaryFirst step in NS-Forest v4.0

The BinaryFirst step is introduced in version 4.0 of NS-Forest and is implemented in the gene pre-selection step of the workflow. Essentially, this process reduces the number of genes that are later considered in the random forest step as candidate marker genes for each cluster by only selecting genes with a Binary Expression Score greater than or equal to a dataset-specific threshold based on the distribution of this score (Figure 2). The Binary Expression Scores are first calculated for each gene-cluster pair in the dataset using the formula defined in the paper detailing version 2.0^13^. Users can specify which type of threshold is used in the BinaryFirst step: ‘none’, ‘BinaryFirst_mild’, ‘BinaryFirst_moderate’, or ‘BinaryFirst_high’. If the threshold is ‘none,’ no filtering is performed, and all genes in the input anndata object are considered for the iterative search in the random forest step. The other threshold values are calculated based on the distribution of Binary Expression Scores from all genes, and so these values vary depending on the input dataset used. The mild threshold is set as the median Binary Expression Score, the moderate threshold is set as the mean Binary Expression Score plus the standard deviation of all scores, and the high threshold is set as the mean Binary Expression Score plus two times the standard deviation. In version 4.0, the default is set to ‘BinaryFirst_high,’ which is the most stringent threshold that filters the most genes.

The median Binary Expression Score in a scRNA-seq dataset is often 0, as is the case for the human MTG, kidney, and lung datasets used in this paper. This is expected because most genes are not useful for distinguishing granular cell types at single cell resolution. In such cases, the ‘BinaryFirst_mild’ model in version 4.0 is equivalent to the model used in version 2.0/3.9, since no initial filtering is done when NS-Forest is run with this threshold. Thus, the results obtained from running version 4.0 with the ‘BinaryFirst_mild’ threshold is equivalent to results obtained from running NS-Forest version 2.0 or 3.9 (no algorithmic difference between versions 2.0 and 3.9). By default, NS-Forest v4.0 uses the ‘BinaryFirst_high’ threshold, unless otherwise stated.

For the human middle temporal gyrus (MTG) dataset, the median Binary Expression Score is 0, indicating that more than half of the original input genes have zero median expression in these clusters and are therefore non-informative. In the human MTG dataset, the mean Binary Expression Score is 0.167 and the standard deviation is 0.249, implying the thresholds for this specific dataset are as follows: mild = 0, moderate = 0.176 + 0.249 = 0.415, high = 0.176 + 2*0.249 = 0.664. For the human kidney and lung datasets, the median Binary Expression Score is also 0. The moderate and high thresholds are 0.102 and 0.194 for the kidney dataset, respectively, and 0.118 and 0.222 for the lung dataset, respectively.

All runtimes discussed in the Results section detailing the improvement obtained with the BinaryFirst step were obtained from running NS-Forest through jobs that were submitted to the *Expanse* supercomputer system in the San Diego SuperComputer Center.

### On-Target Fraction metric

In version 4.0, a new On-Target Fraction metric is provided, to quantify the expression specificity of the marker genes with respect to their target cell types. In previous versions, the algorithm reported the F-beta score and Positive Predictive Value (PPV) together with the true/false positives/negatives (i.e., TP, FP, TN, FN), which are metrics that quantify the discriminative power of each set of markers for their cell type classification performance. However, these metrics do not fully capture how well NS-Forest achieves the ideal scenario of identifying markers for each cluster that are exclusively expressed in that cluster. To make a clear distinction, we refer to F-beta score and PPV as *classification* metrics, and On-Target Fraction as an *expression* metric.

The On-Target Fraction is defined for each marker gene g in target cluster T as,

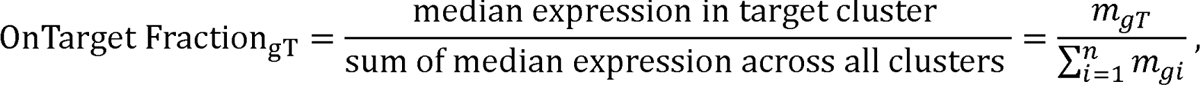

where m_gT_ = median expression of marker gene g in target cluster T, and m_gi_ = median expression of marker gene g in cluster i for i = 1, …, n. This metric has a range of [0,1], with a value of 1 being the ideal case where this marker gene is “exclusively” expressed in more than half of the cells in its target cluster and in fewer than half of the cells in all other clusters. Due to the zero-inflation nature of scRNA-seq data, this metric can effectively capture those genes that have abundant expression exclusively within the target cluster. At the cluster level, we report the On-Target Fraction for each cluster using the median Fraction_gT_ of its marker genes. We used the median as a summary statistic when reporting the On-Target Fraction to account for the non-normal nature of scRNA-seq gene expression distribution.

### NS-Forest Python package

With the continuous refinements of the NS-Forest algorithm and its marker evaluation metrics, NS-Forest has become a comprehensive software package. To provide a user-friendly software package, NS-Forest v4.0 is now modularized, consisting of 4 main functional modules: preprocessing, NSForesting, evaluating, and plotting. NS-Forest v4.0 takes in an anndata^15^ object in .h5ad format that contains a cell-by-gene expression matrix and the cell type cluster membership column stored in the observation-level (.obs) metadata matrix of the data object.

An optional but suggested step before preprocessing is to generate a hierarchical clustering dendrogram of the cell type clusters on the full dataset. This occurs before any gene filtering because the dendrogram should be consistent between various preprocessing methods. In the preprocessing module, the first step is to calculate the median expression matrix for each gene in each cluster. The default positive_genes_only parameter is true, which filters for genes with a positive median expression in at least one cluster. (Note that this preprocessing step based on medians is specific to NS-Forest and should not be used if evaluating the performance of marker genes produced by other approaches.) Based on the median expression matrix, the Binary Expression Score of each gene (positive genes only by default) is calculated in each cluster. The Binary Expression Score has a range of [0, 1], with values closer to 1 indicating a higher level of binary expression (i.e., the gene is expressed in the target cluster and not others). The pre-calculated median expression matrix and the Binary Expression Score matrix are saved in the unstructured metadata slot (.uns) of the data object.

In the NSForesting module, the BinaryFirst step is implemented with the gene_selection parameter that determines the BinaryFirst criterion (None: 0, BinaryFirst_mild: median, BinaryFirst_moderate: mean + std, and BinaryFirst_high: mean + 2std), which filters out genes below the chosen threshold. The main step in the NS-Forest algorithm is building a random forest classifier for each cluster that is trained on the genes that passed the gene_selection criteria. In this step, each gene is ranked by the Gini Impurity index and the n_top_genes with the highest Gini index are then reranked by their pre-calculated Binary Expression Scores. The n_gene_eval value indicates how many genes from the reranked candidate gene list are input into the decision tree evaluation for determining the best combinatorial marker genes as final output. A single split decision tree is built for each evaluated gene. Gene combinations of all set lengths are evaluated, and the combination with the highest F-beta score is considered the final set of NS-Forest marker genes for that cluster. The performance metrics returned from this module for each cluster are the F-beta score, PPV (precision), recall, TP, FP, TN, FN, and On-Target Fraction.

The evaluating module can be called independently without calling the preprocessing and NSForesting module. It is useful for calculating the metrics for a user-input marker gene list with paired cluster names to compare the cell type classification performance and marker expression across different marker gene lists. We provide an option of using mean instead of median for the On-Target Fraction calculation, to account for cases where a user-input gene is only expressed in a small proportion of cells in the target cluster and has absolutely no expression in other clusters.

The plotting module creates the scanpy^15^ dot plot, stacked violin plot, and matrix plot figures for visualization of the NS-Forest or user-input marker genes with clusters organized according to the dendrogram order (from the preprocessing step) or in order corresponding to user-input. Other plotting functions include creating interactive plotly^25^ boxplots and scatter plots, which are useful for comparing metrics and identifying clusters of interest.

## Data Availability

The human brain Middle Temporal Gyrus (MTG) dataset was downloaded from https://portal.brain-map.org/atlases-and-data/rnaseq/human-mtg-smart-seq. The human kidney dataset was downloaded from https://cellxgene.cziscience.com/collections/bcb61471-2a44-4d00-a0af-ff085512674c (Integrated Single-nucleus and Single-cell RNA-seq of the Adult Human Kidney). The human lung dataset is currently unpublished (manuscript under submission). The Human Lung Cell Atlas (HLCA) dataset was downloaded from https://cellxgene.cziscience.com/collections/6f6d381a-7701-4781-935c-db10d30de293 (core).

## Code Availability

NS-Forest v4.0 is openly available at https://github.com/JCVenterInstitute/NSForest, with documentations and tutorials available at https://nsforest.readthedocs.io.

## Acknowledgments

This work was supported by the U.S. National Institutes of Health (1RF1MH123220, 1R03OD036499, OT2OD033756, U54HL165443, and U54HL145608) and the Intramural Research Program of the National Library of Medicine (NLM), National Institutes of Health. The funding bodies had no role in the design or conclusions of this study.

## Author contributions

A.L., R.H.S. and Y.Z. conceived the project. A.L. and Y.Z. designed and implemented the algorithm. B.P. built the software package. A.L., B.P. and A.V.P. conducted the data analyses. T.E.D. and G.P. provided the unpublished data and supervised the analysis on it. A.L., B.P., R.H.S., and Y.Z. interpreted the results. A.L., B.P., R.H.S., and Y.Z. wrote the manuscript. All authors agreed on the contents of the manuscript.

---

**Supplementary Figure 1.**
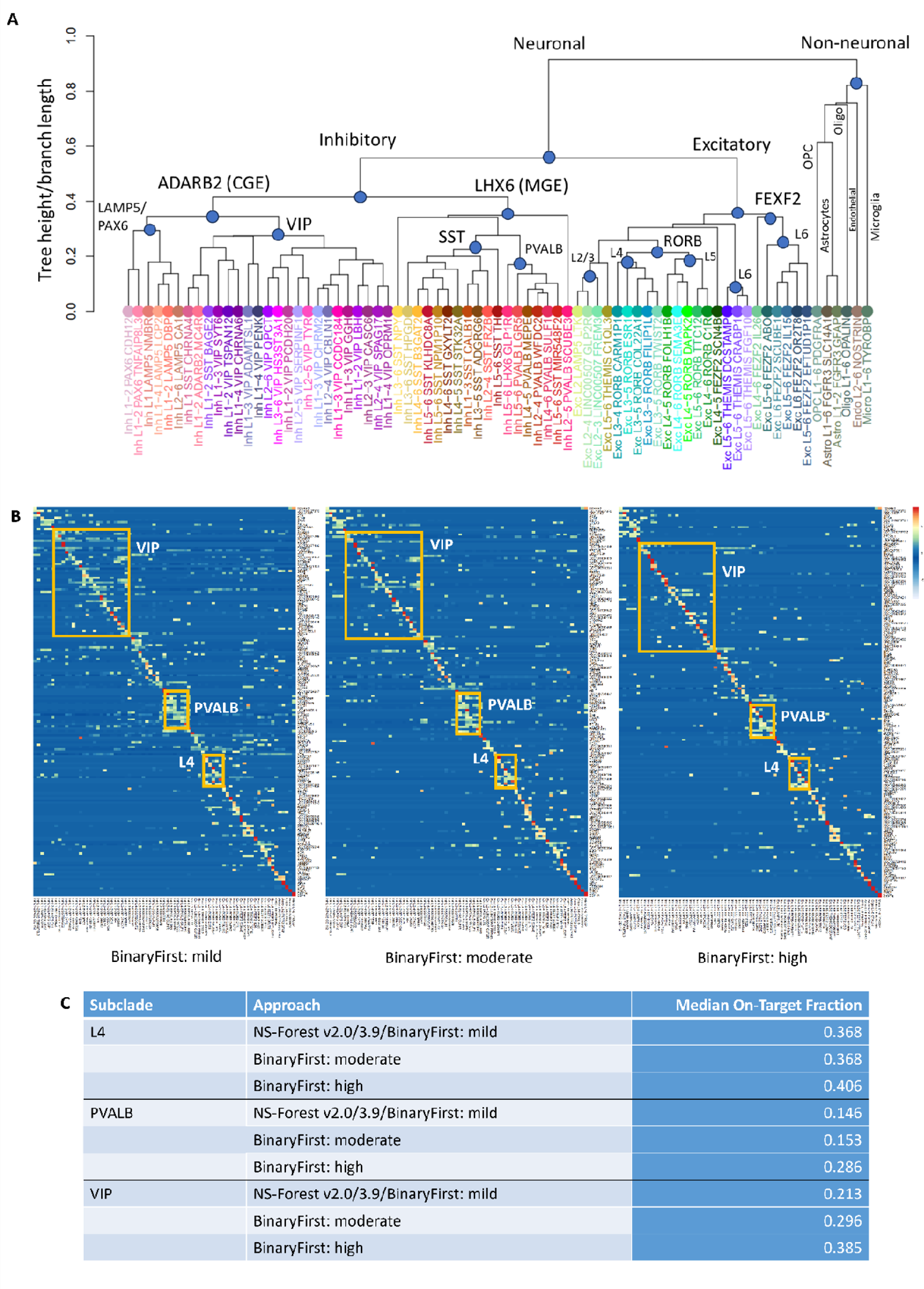
Comparing performance of using different BinaryFirst thresholds in NS-Forest v4.0 on specific subclades within human MTG dataset. (A) Hierarchical dendrogram derived in the original human MTG study with labelled and color-coded subclades (https://github.com/AllenInstitute/MOp_taxonomies_ontology/tree/main). (B) Heatmaps of markers from the human middle temporal gyrus (MTG) dataset generated from NS-Forest v4.0 with ‘BinaryFirst_mild’, ‘BinaryFirst_moderate’, and v4.0 with the ‘BinaryFirst_high’ thresholds. The regions on the heatmaps highlighted by the orange boxes correspond to the identified markers for the cell types in specific subclades (VIP, PVALB, and L4 subclades) that are known to be more similar to each other and thus, more difficult to distinguish. The colors correspond to the normalized median expression level (log2-transformed counts per million) for the marker gene (rows) in a given cell type cluster (columns), with high expression in red/yellow, and low expression in blue/white. The clusters are ordered according to the hierarchical dendrogram provided in the original study shown in (A). (C) Median On-Target Fraction values within each of the three specific subclades across these three BinaryFirst thresholds.

**Supplementary Figure 2.**
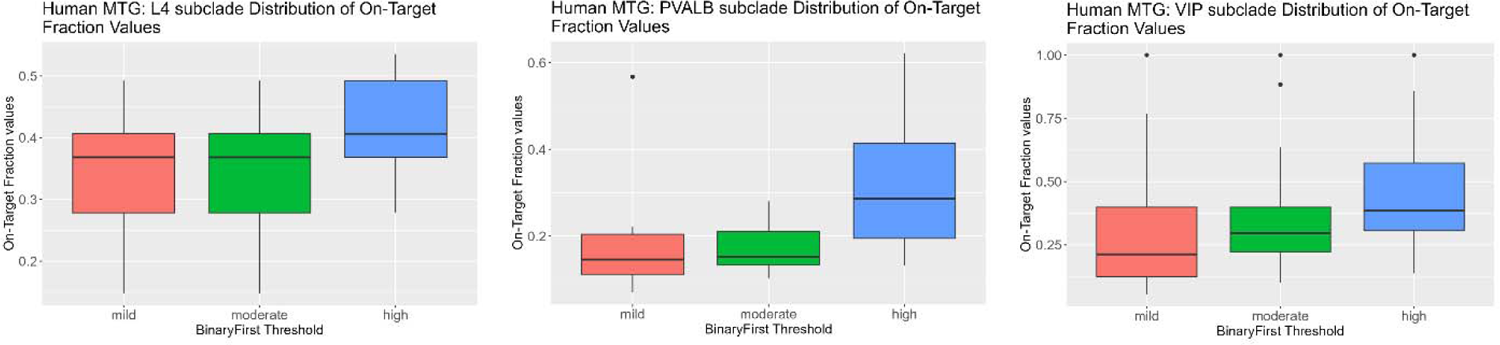
Comparison of On-Target Fraction distribution in major subclades of Human MTG dataset across BinaryFirst thresholds. Boxplots comparing the distribution of the On-Target Fraction values within each of the three specific subclades in the human MTG dataset (L4, PVALB, and VIP subclades) across the mild, moderate, and high BinaryFirst thresholds are shown.

**Supplementary Figure 3.**
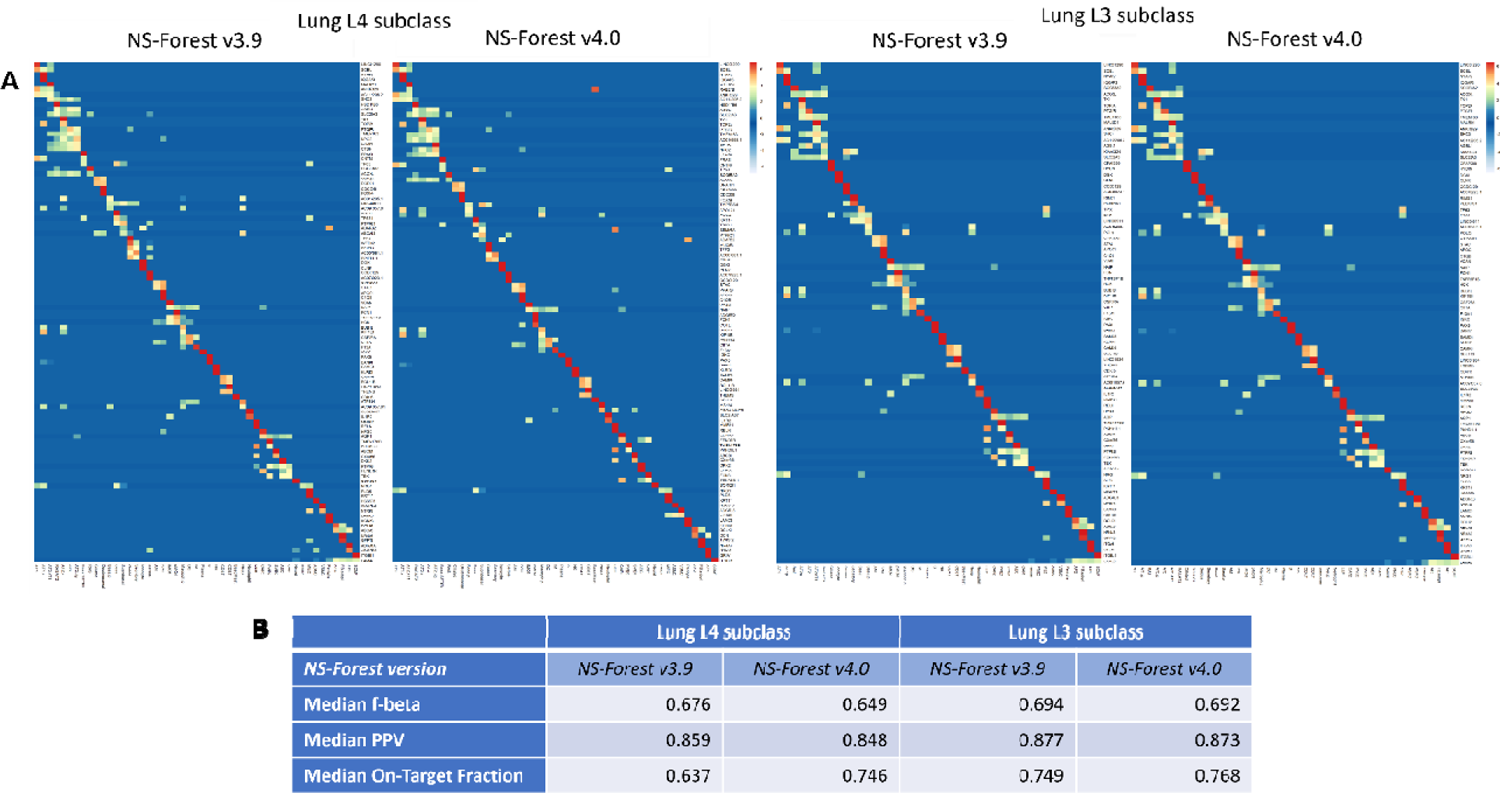
Evaluating performance of NS-Forest on lung L4 and L3 subclass datasets. (A) Heatmaps of NS-Forest v2.0/v3.9 and v4.0 markers from the L4 and L3 subclasses of the human lung. The colors correspond to the normalized median expression level (log2-transformed counts per million) for the marker gene (rows) in a given cell type cluster (columns), with high expression in red/yellow, and low expression in blue/white. The clusters are ordered according to the hierarchical ordering in the dendrogram generated by the scanpy package (*scanpy.tl.dendrogram*) using default settings. **(B)** Comparison of the performance metrics corresponding to the NS-Forest results shown in (A).

**Supplementary Figure 4.**
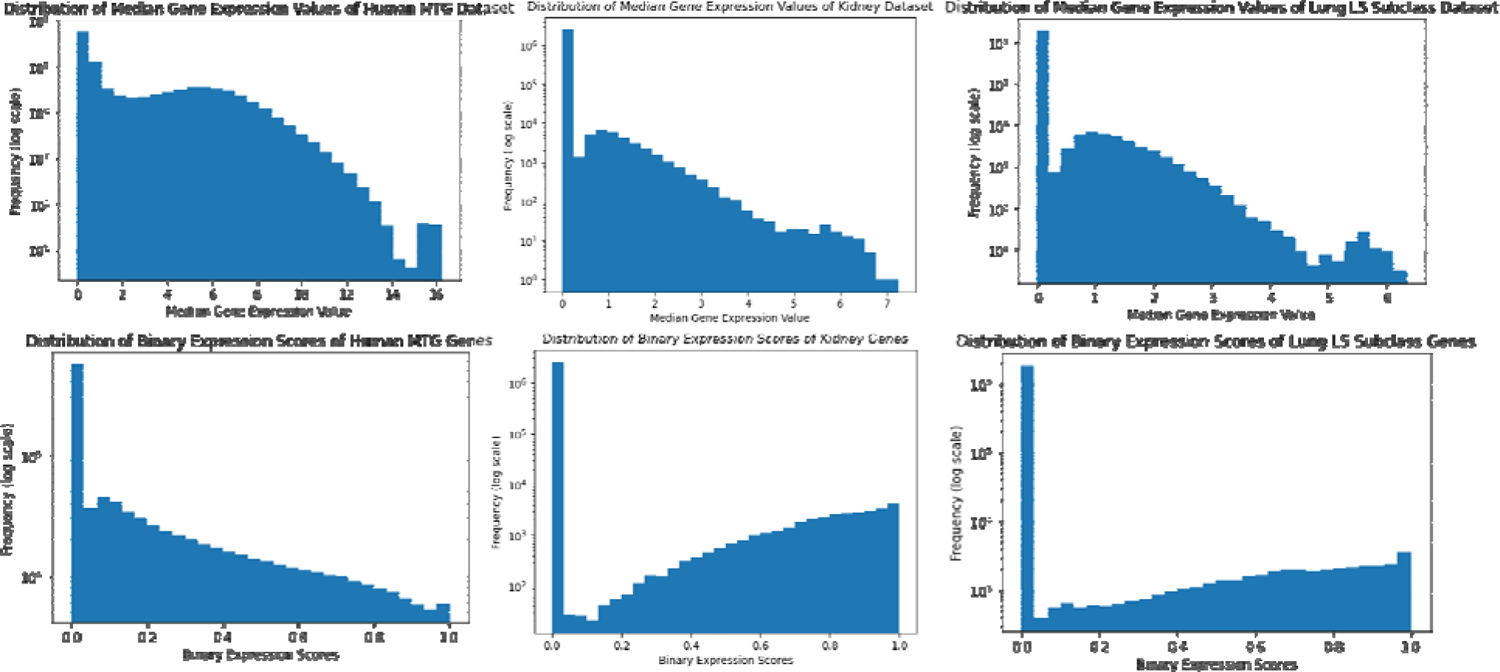
Distribution of median gene expression per cluster and Binary Expression Score in human MTG, kidney, and lung datasets. First row: histograms of distribution of median gene expression values of genes expressed in all clusters in human MTG, kidney, and lung datasets. X-axis displays the range of median gene expression values in each dataset, and the y-axis displays the frequency of each median gene expression value (log scale). Second row: histograms of distribution of Binary Expression Score values of genes in these three datasets. X-axis ranges from 0 to 1 (representing the possible values the binary expression score can be), and the y-axis displays the frequency of each binary score value (log scale). All distributions are highly right-skewed.

**Supplementary Figure 5.**
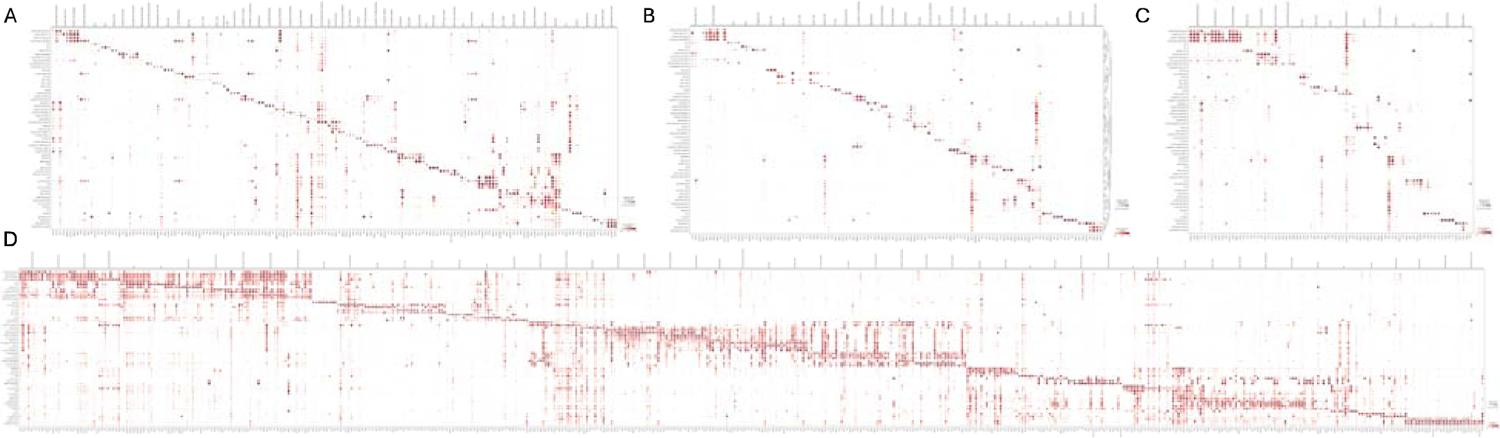
Dotplots of HLCA, CellRef, ASCT+B, and Azimuth marker genes on HLCA core cell types. (A) 162 HLCA markers across 61 HLCA cell types. **(B)** 115 CellRef markers across 33 HLCA cell types. **(C)** 80 ASCT+B markers across 18 HLCA cell types. **(D)** 535 Azimuth markers across 56 HLCA cell types.

**Supplementary Figure 6.**
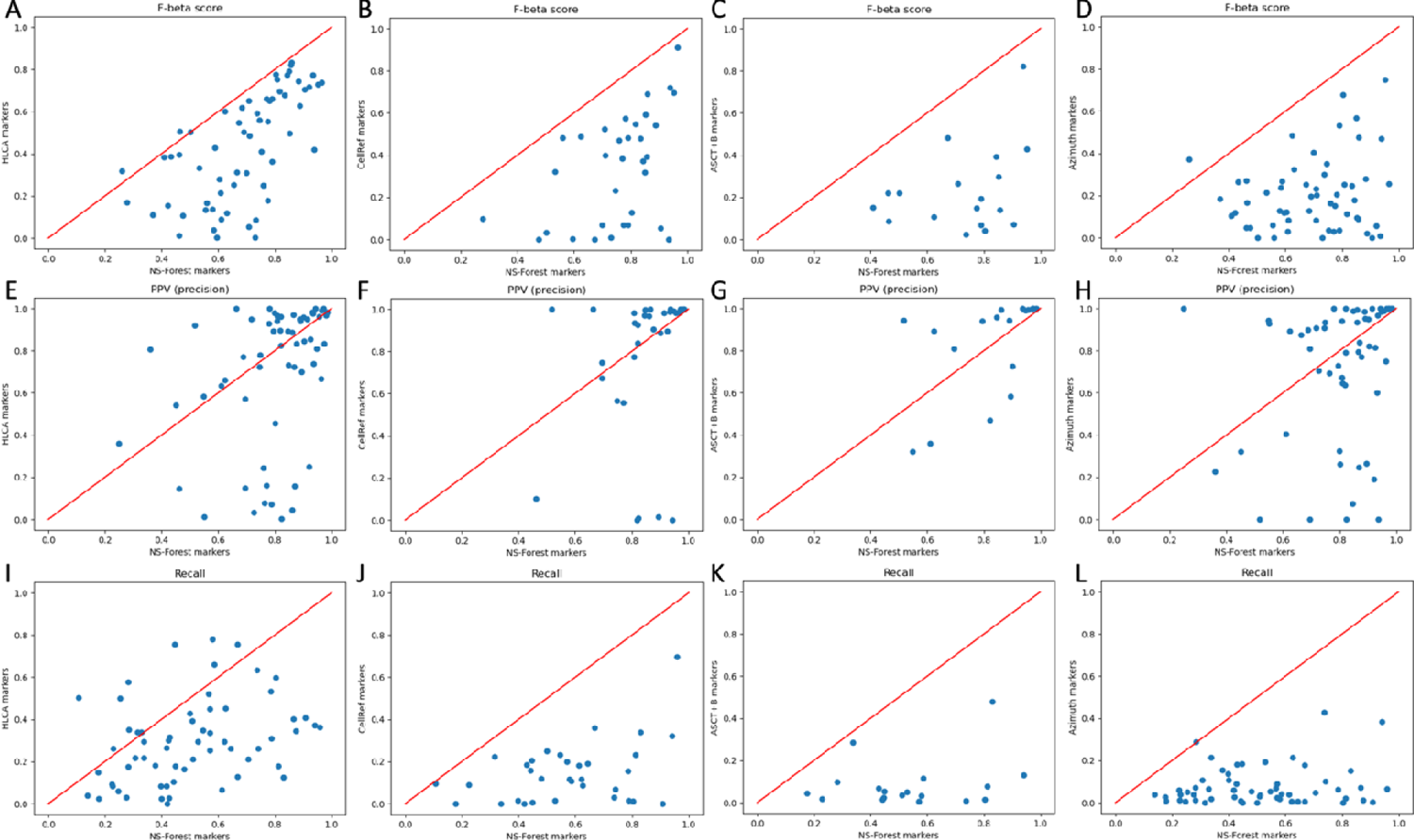
NS-Forest consistently outperforms other published methods in classification performance. (A-D) Scatter plots comparing F-beta scores for each cell type using NS-Forest markers vs. HLCA, CellRef, ASCT+B, and Azimuth markers. **(E-H)** Scatter plots comparing PPV (precision) for each cell type using NS-Forest markers vs. HLCA, CellRef, ASCT+B, and Azimuth markers. **(I-L)** Scatter plots comparing recall for each cell type using NS-Forest markers vs. HLCA, CellRef, ASCT+B, and Azimuth markers.

**Supplementary Table 1.** NS-Forest v4.0 results of the human brain middle temporal gyrus dataset.

**Supplementary Table 2.** NS-Forest v4.0 results of the human kidney dataset.

**Supplementary Table 3.** NS-Forest v4.0 results of the human lung L5 subclass dataset.

**Supplementary Table 4.** NS-Forest v4.0 results of the human lung L4 subclass dataset.

**Supplementary Table 5.** NS-Forest v4.0 results of the human lung L3 subclass dataset.

**Supplementary Table 6.** NS-Forest v4.0 results of the Human Lung Cell Atlas dataset.

