## Supplementary figures and images for "Discovery of optimal cell type classification marker genes from single cell RNA sequencing data"

### newsuppfig1.png

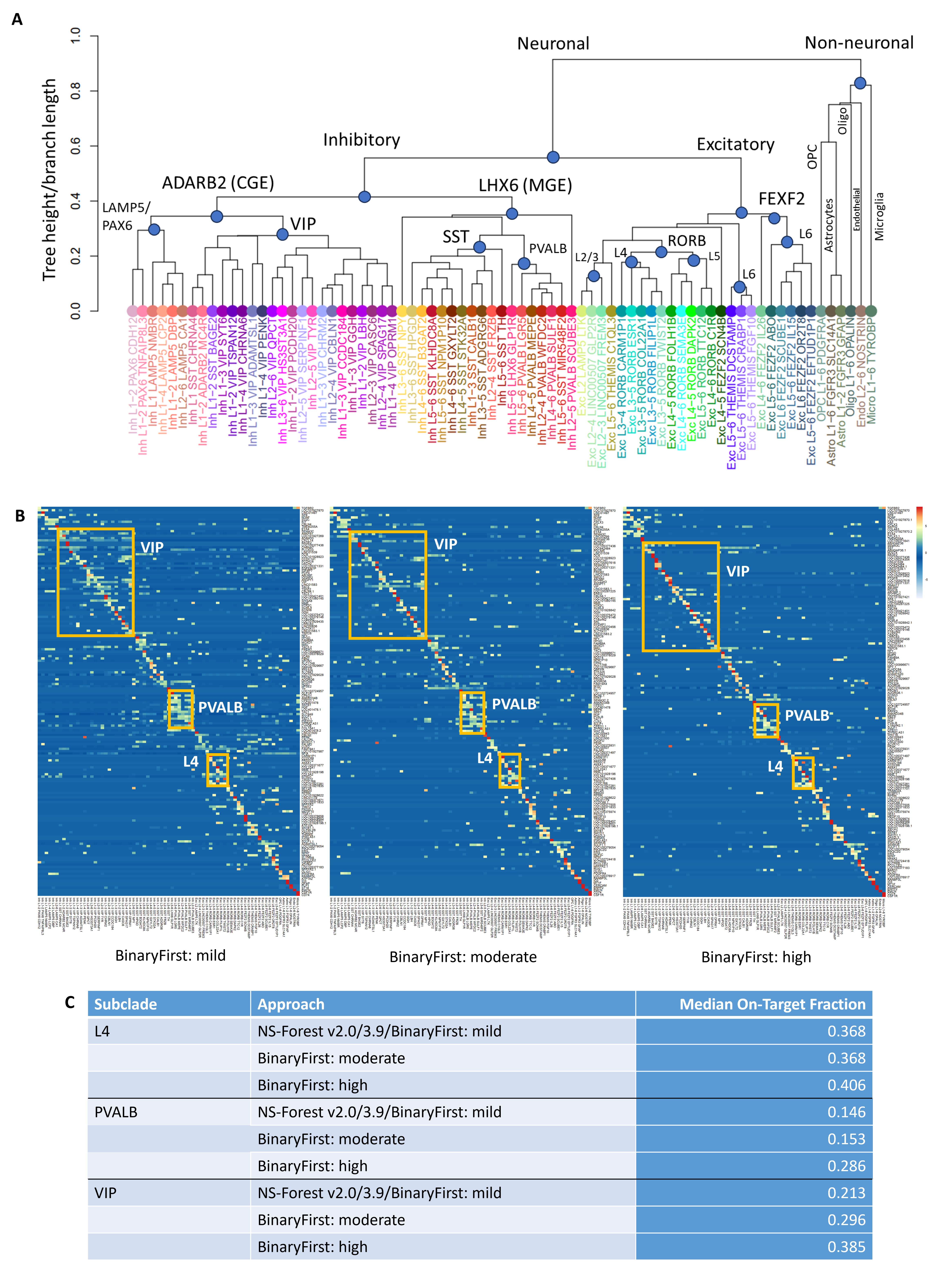

### newsuppfig2.png

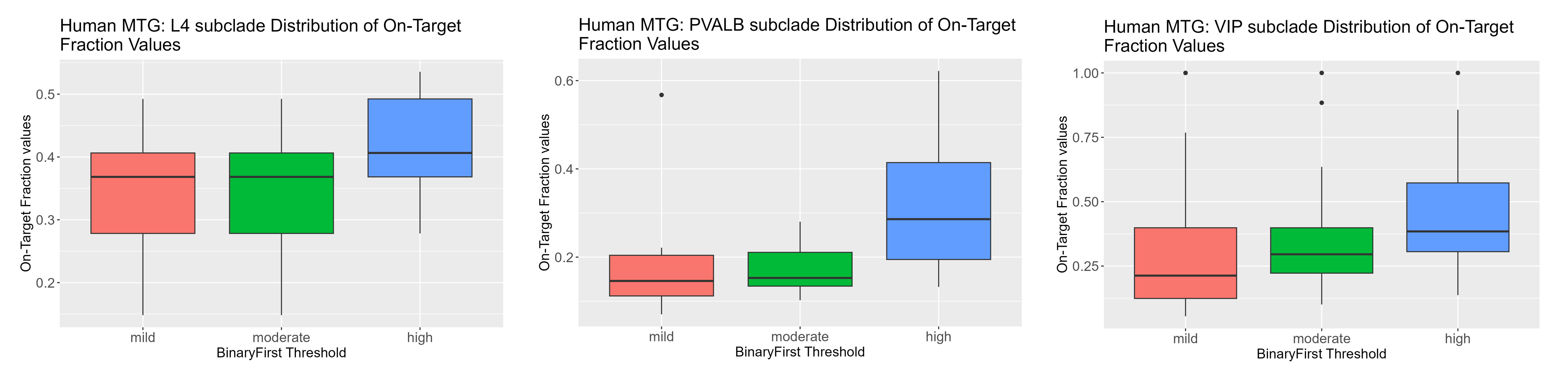

### suppfig3.png

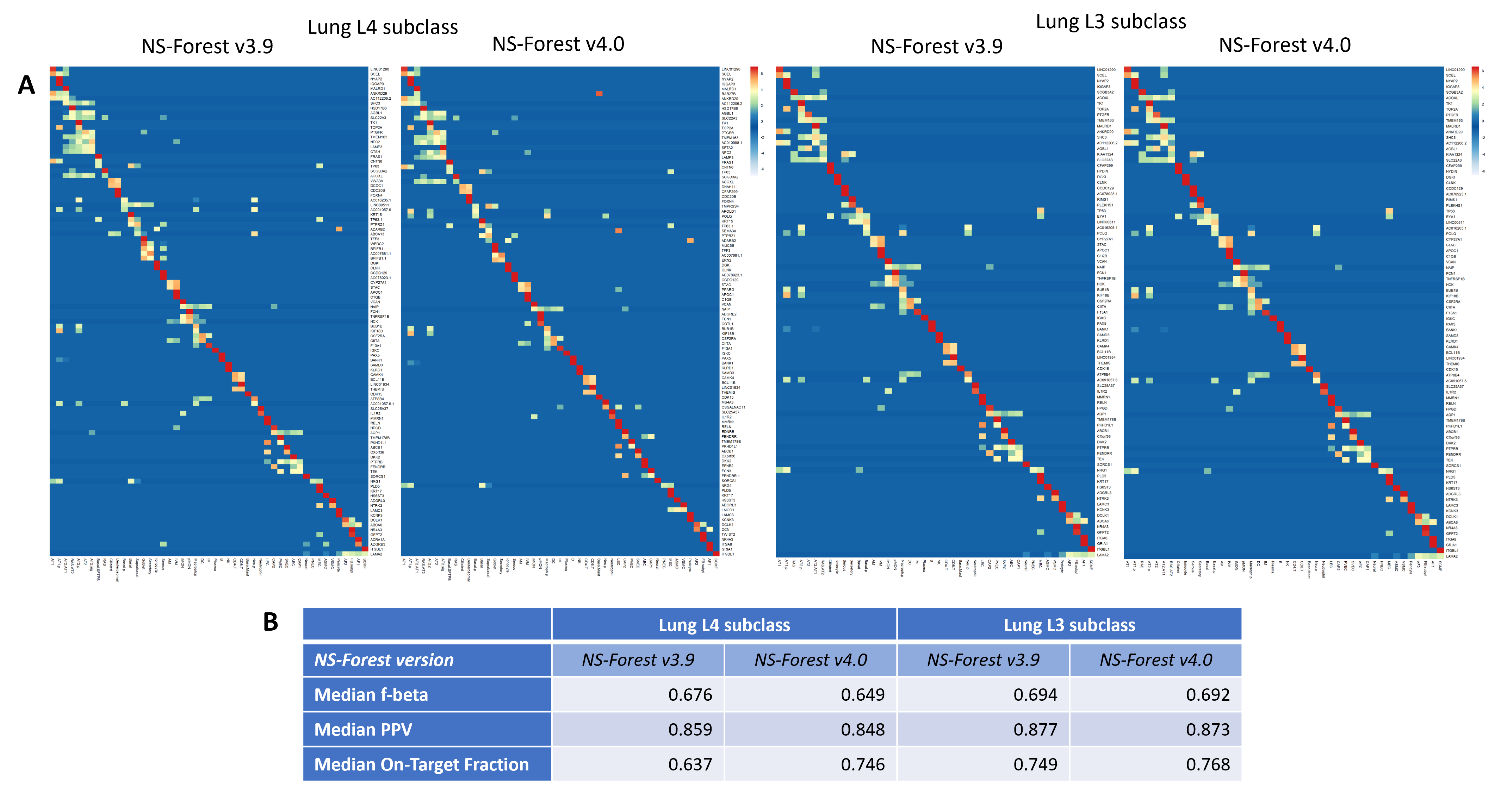

### suppfig4.png

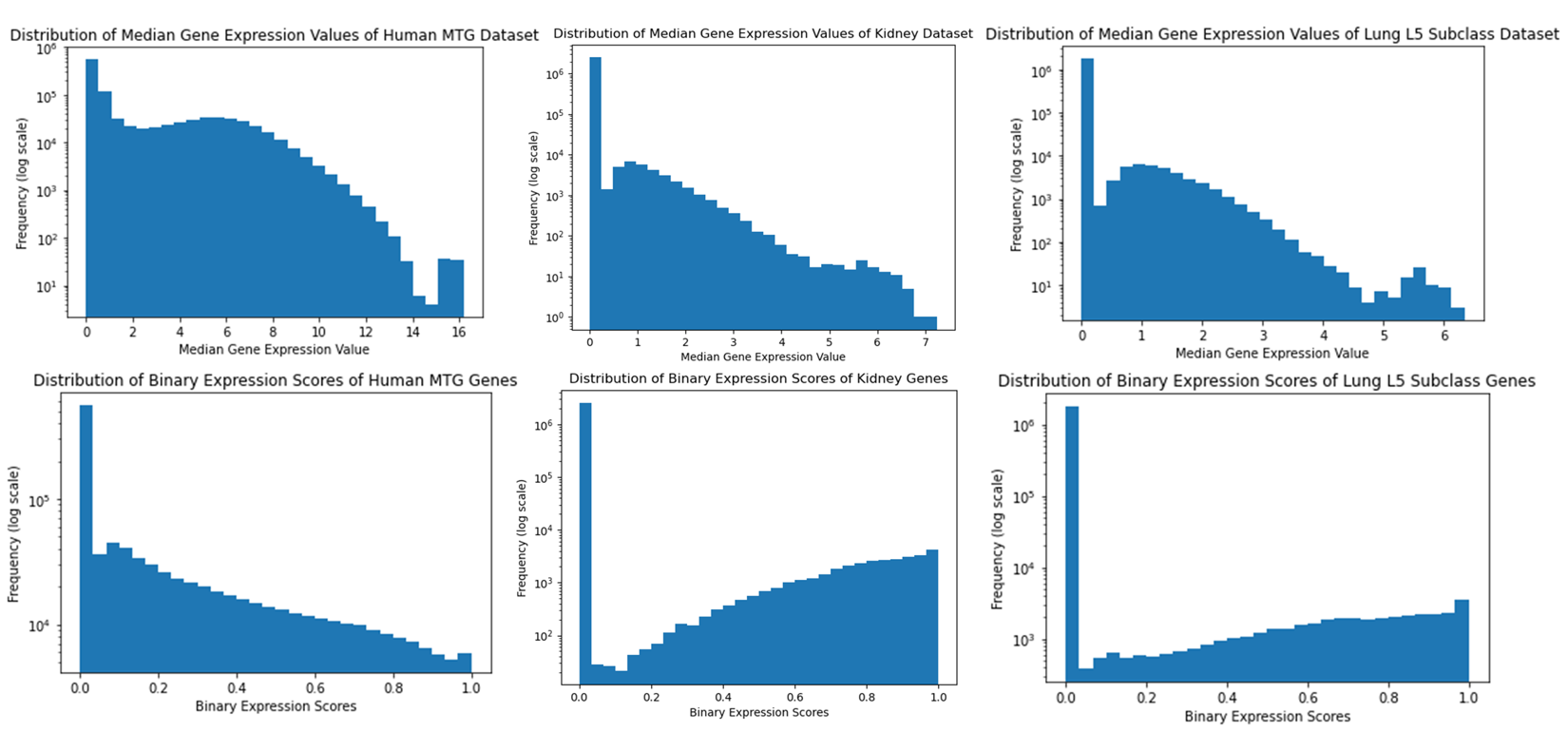

### suppfig5.pdf

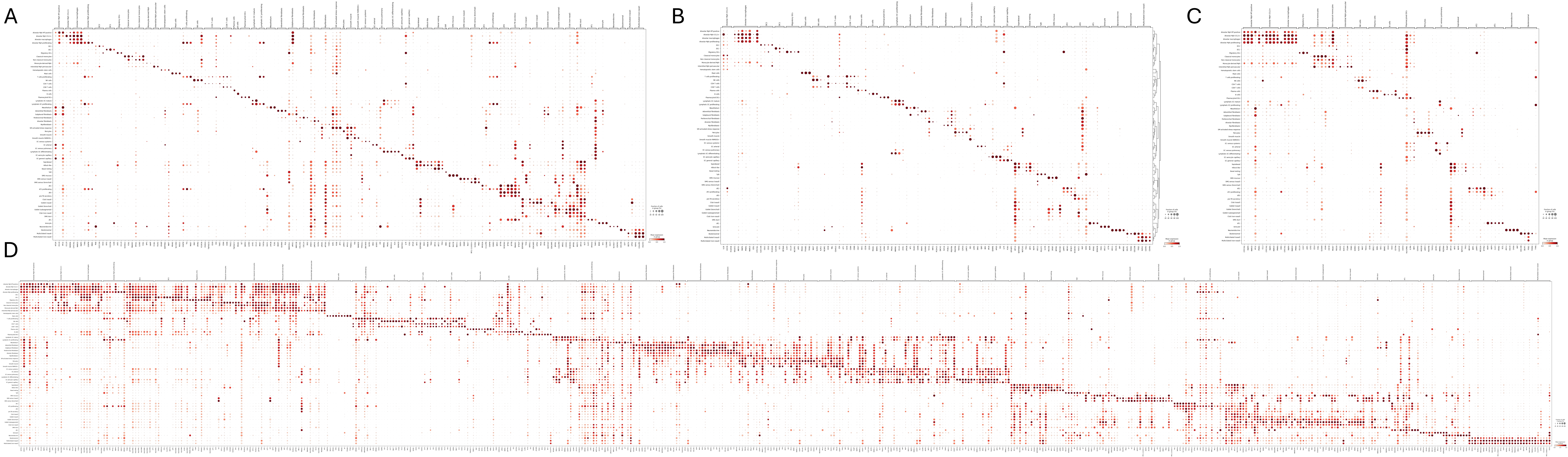

### suppfig6.png

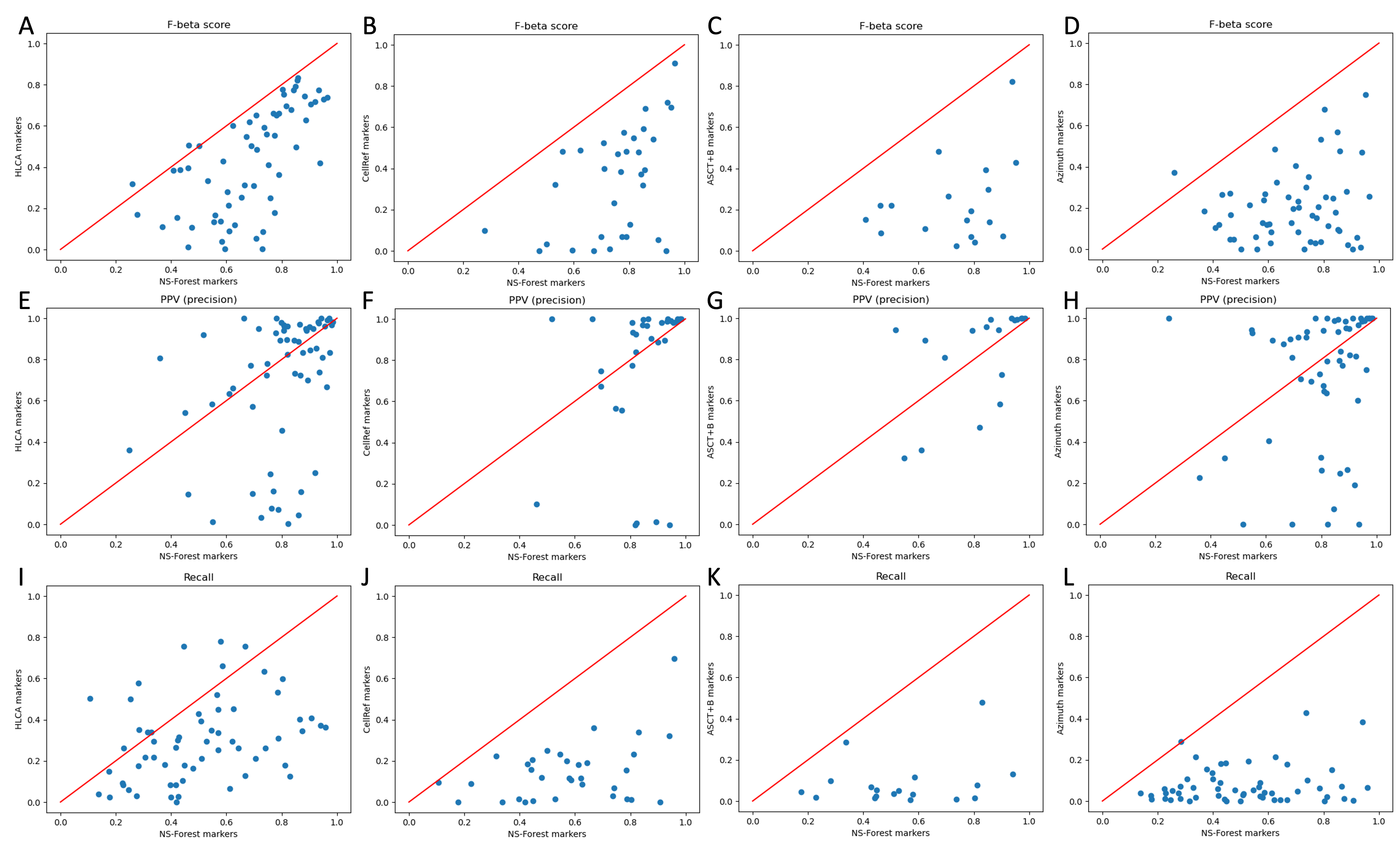
